# Augmented Feedback Training for Overcoming the Learning Plateau of Motor Expertise

**DOI:** 10.1101/2025.10.23.683818

**Authors:** Takanori Oku, Masato Hirano, Kengo Matsuzaka, Shinichi Furuya

## Abstract

Skilled performers often encounter a plateau in which further practice yields little improvement of intricate sensorimotor skills. Overcoming such limits requires novel training paradigms that can engage performers in novel ways of exploring and refining their actions. Here, we introduce a training pipeline that integrates high-precision motion sensing with augmented feedback to enhance expert-level motor learning. Using piano performance as a model, trained pianists practiced imitating a prize-winning expertʼs performance of a technically demanding movement sequence. While conventional auditory-based learning offered limited benefit, augmenting practice with trial-by-trial visualizations of discrepancies between the pianistʼs own movements and those of the expert enabled learners to refine their performance. This training induced richer movement exploration, facilitated closer convergence toward expert motion patterns, and produced perceptible improvements in sound quality evaluated by expert pianists. These findings demonstrate that augmented feedback can break performance plateaus in experts by providing externally-sensed, task-specific information that expands exploration beyond habitual training strategies. Such augmented learning promises new applications across domains where expertise is constrained by entrenched motor habits, including musical performance, athletics, and surgical training.

## Introduction

Recent advances of sensing technologies have transformed the landscape of motor skill learning by enabling augmented feedback, which provides supplementary information beyond what the human sensory system can naturally perceive. Such feedback has emerged as a powerful means of refining complex skills by making otherwise imperceptible features of movement accessible for conscious correction and update of the skills. This approach is particularly valuable for domains requiring millisecond-level timing and fine-grained spatial control, such as sports, microsurgery, and music performance (Grosshauser & Hermann, 2009; Hicheur et al., 2020a; Lee & Schmidt, 2014; Oku & Furuya, 2019; Sigrist et al., 2013; Takaishi et al., 2023; Van Der Meijden & Schijven, 2009). By mapping expert movements or ideal movement trajectories onto visual, auditory, or haptic modalities, augmented feedback creates a training pipeline in which learners iteratively compare their own actions with expert benchmarks and imitate adaptively (Gorman et al., 2022; Todorov et al., 1997). This form of imitation-based training can facilitate the acquisition of novel skills and enhance accuracy in novices who have limited prior knowledge of the ideal task performance and task itself (Hanashima et al., 2023; Takahashi et al., 2019). However, despite its promise, most of previous applications of the augmented feedback have primarily targeted beginners and intermediate learners (Petancevski et al., 2022), which limits the understanding of its efficacy for breaking through the entrenched plateaus of trained individuals. Addressing this gap is essential, not only for advancing theories of motor learning but also for designing novel training systems that integrate human adaptability with machine precision.

For experts of motor tasks, simple augmented feedback training often proves ineffective or yields only limited benefits (Petancevski et al., 2022). One reason for this limited effectiveness is that experts possess superior sensory and motor control abilities specific to the task (Hirano, Sakurada, et al., 2020; Hosoda & Furuya, 2016), reducing the relative value of externally provided information. Furthermore, frequent feedback can lead to excessive attention to movements, hindering the automatization of skills (Beilock & Gray, 2012). Various methods have been proposed to overcome this issue, such as multimodal feedback to increase informational richness (Abiri et al., 2019; FU et al., 2025; Hicheur et al., 2020b; Pardue & McPherson, 2019; Riley et al., 2005), and selective or reduced feedback frequency (Carter et al., 2014; Chiviacowsky & Wulf, 2002; van der Meer et al., 2024; Wulf, 2007). Nevertheless, the optimal design of augmented feedback tailored for experts remains unclear. Piano performance provides an appropriate testbed for exploring augmented feedback methods effective for experts. This sensorimotor skill requires precise control of timing and force of sequential movement sequences with multiple fingers (Furuya et al., 2011; Oku & Furuya, 2017). Various methods for acquiring the skill of piano performance through augmented feedback have been proposed, although most of these studies target beginners or intermediate players (Hamond et al., 2019; Nusseck et al., 2025; Oku & Furuya, 2019; Riley et al., 2005). Skilled pianists are characterized by sophisticated motor skills such as anticipatory modification of individual movement elements (Winges et al., 2013) and independent movement control of multiple fingers (Kimoto et al., 2022). Considering the specialized motor skills of expert pianists, we hypothesized that augmented feedback training for expert pianists should target not only timing or force errors of individual movement elements, but also high-precision measurements of key movements, which are difficult to be perceived accurately via the sensory receptors.

In this study, we developed a novel augmented feedback training system with the high-precision sensing system of the key movements in order to facilitate imitation training for expert pianists. Imitating expert performers is a challenging motor task due to precision and speed of a succession of the individual movements that occur simultaneously and independently. Nevertheless, as demonstrated by the implicit knowledge transfer from coaches to learners, imitation has been an essential method for expert performers to refine their skills. Specifically, we designed a system that visualizes the differences in time-varying waveforms of key movements between a target performance by an exceptionally-skilled expert pianist who has won international competitions and performances by conservatory-level pianists. We evaluated its effectiveness for skill learning using both objective similarity indices of key motion waveforms and auditory experiments assessing the perceived similarity of sound impressions. Through this approach, we aim to propose a new feedback framework that supports continued skill development of expert pianists.

## Methods

### Experiment 1: Changes in Similarity of Key Movement Patterns through Augmented

#### Feedback Training

This study investigated the effects of the training that provide high-precision feedback on the key movement during piano performance. The time-varying data of the piano key movements were recorded by using the high-resolution sensing system of the piano keys (Oku & Furuya, 2022). The sensor optically measures the displacement of all 88 keys of the piano with the spatial resolution of 0.01 mm and temporal resolution of 1 ms in a non-contact manner. The spatiotemporal resolution of the sensing system is superior to that of human intrinsic sensor systems (Borngräber et al., 2022; Friberg & Sundberg, 1995; Van Boven & Johnson, 1994). The recorded key movement data were visually displayed and compared with the reference performance recorded from an expert pianist who won prizes at international competitions for professional pianists. Figure 1 illustrates the snapshot of the visual feedback provided for a participant, in which solid and dashed lines represent the key movements of the participant and the target performance.

**Figure 1.**
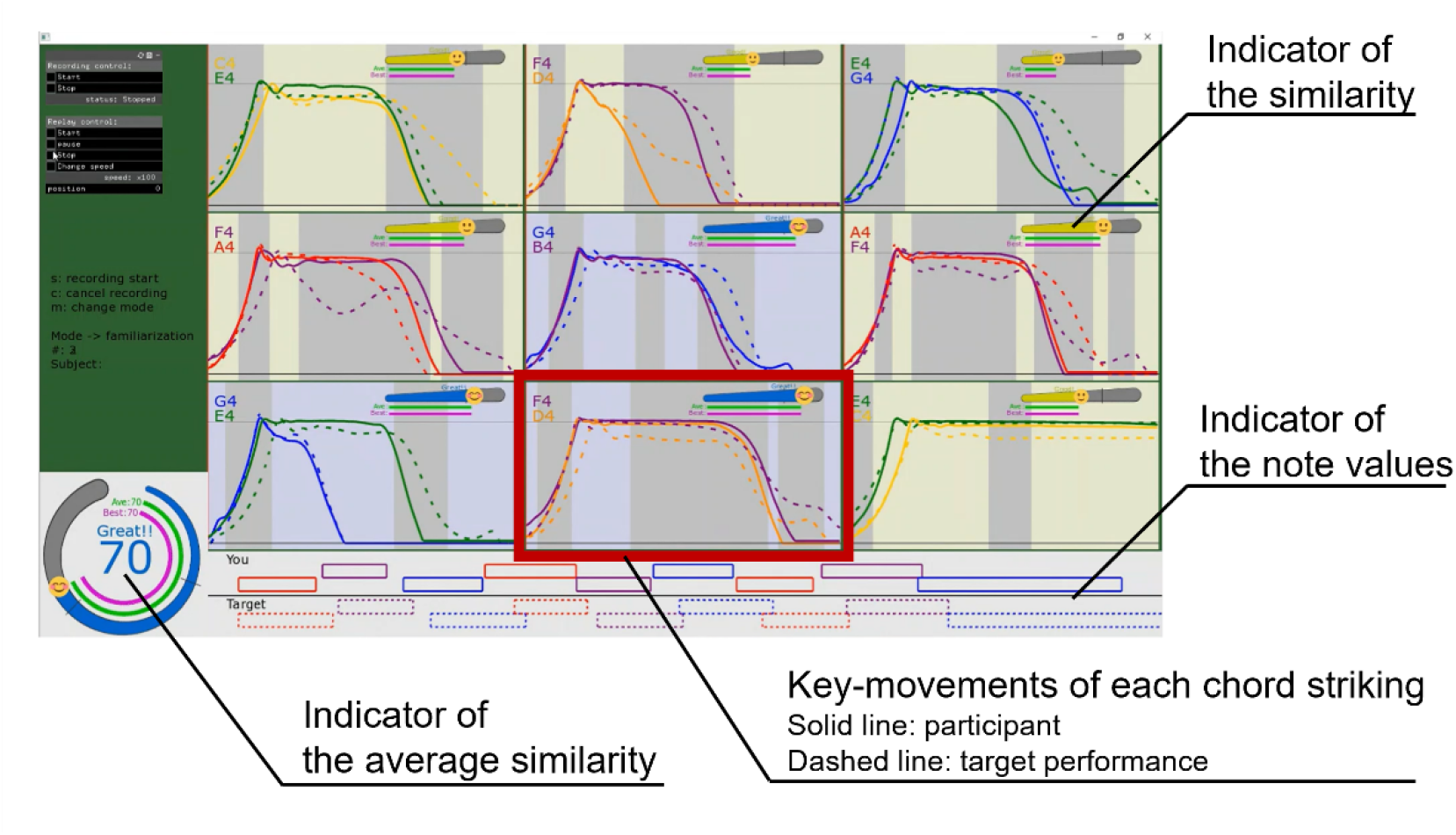
A snapshot of the visual feedback provided during the augmented FB training session. Piano key movements during performances were recorded using a custom-designed sensing system with high spatiotemporal resolutions. The visual feedback was displayed on a monitor put beside the participant. The solid and dashed lines represent the key movements extracted from the latest trial by the participant, and the target key movements, respectively. Different colors in line indicate movements of different keys. The top right indicator in each window represents the similarity score of the key movements between the participant and the target performance for each of the nine chords. The lower left indicator displays the average similarity across all chords, providing a measure of similarity of the overall performance. At the bottom, the duration of the key depression (i.e. note value) for each chord was displayed for the participant (top) and target performance (bottom).

Similarity of the key movements between the participant and the reference performance was evaluated by computing the cosine similarity of the two key waveforms for each chord. The similarity of a chord was defined as the mean cosine similarity across all notes constituting the chord. To compute cosine similarity, the waveforms of each keystroke from both the reference (expert) and the participant were temporally aligned. Since the calculation of cosine similarity requires equal length vectors, the time window for each chord comparison was defined as the longer chord-striking duration between the target and the participant. Here, the chord-striking duration refers to the interval from the onset of the first note to the offset of the last note within the chord. The start of the alignment window was set to the onset of the first keystroke in each chord. For instance, in the target performance, if two notes constituting a chord (A and B) were struck in the sequence A‒B at quite short interval, whereas a participant struck the corresponding notes in the sequence B‒A, the time-series waveform of the key movement for the participant’s note A was extracted starting from the timing of the keystroke on note B. This alignment procedure was designed to preserve the temporal characteristics of chord execution when calculating similarity. The temporal discrepancies and imbalances in the durations of notes had a detrimental effect on the similarity calculation with the alignment algorithm. The displayed similarity score was defined as follows.

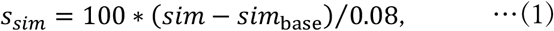

where *s*sim, *sim*, and *sim*base represent similarity score, the cosine similarity between the participantʼs performance and the target, and baseline similarity, respectively. The baseline similarity was set to 0.92, which was sufficiently lower than the minimum similarity value observed in a pilot experiment that was performed with six pianists.

#### Participants

In total, 32 participants were randomly assigned to two groups (i.e. 16 per group), each group receiving a different feedback modality during the training sessions. All of the participants were pianists without any history of neurological disorders. All of the pianists majored in piano performance at a musical conservatory. In accordance with the Declaration of Helsinki, the experimental procedures were explained to all participants. Informed consent was obtained from all participants before participating in the experiment, and the experimental protocol was approved by the ethics committee of Sony Corporation (#22-18-0001).

#### Experimental Task

The experimental task consisted of nine sequential triad chord production involving a thumb-under maneuver (Figure. 2A). This type of chord sequence frequently appears in various piano pieces (e.g. Chopinʼs Etude Op. 25, No. 6). Playing this passage with legato has been considered for pianists to be one of the most technically challenging skills. As the target performance, a professional pianist who had won prizes at international competitions for professional pianists was instructed to play the passage as legato as possible at the loudness of forte. The experimental task for the participants was to reproduce this recorded performance, after listening to the recorded sound of the target performance. Both the professional pianistʼs and the participantsʼ performances were recorded under identical conditions; in the same soundproof room, with the same grand piano (KAWAI: SK-2), with the same microphone (NEUMANN: U87 ai), and with the same recording settings.

**Figure 2.**
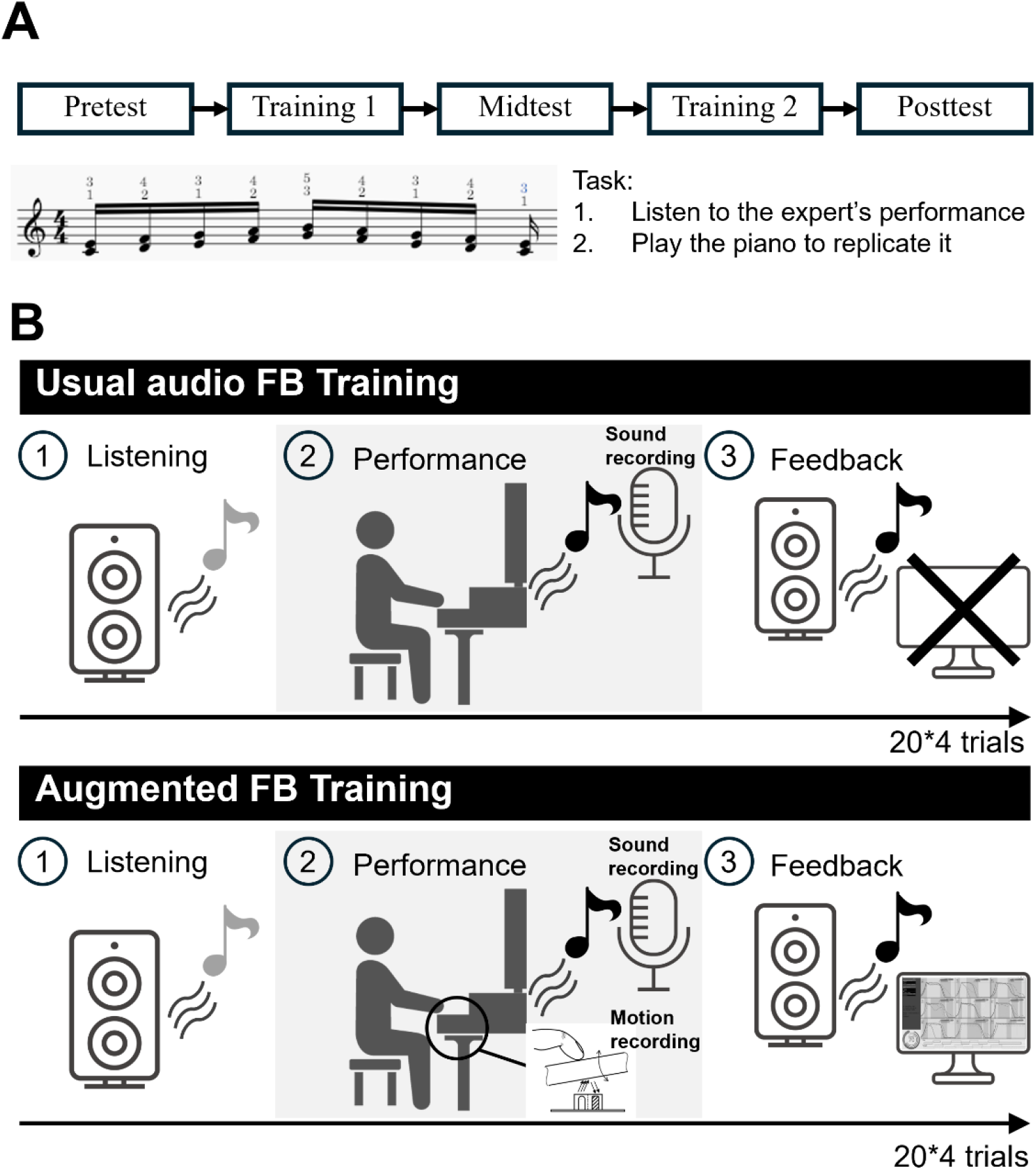
Protocol of the experiment. A: The experiment consisted of three test sessions (Pre-, Mid-, and Post-) with two training sessions between the two of these three test sessions. During the test sessions, participants were instructed to listen to the performance of a highly skilled professional pianistʼs nine sequential triad chord strikings and then reproduce it on the piano. B: In the first training session, the audio FB was administered to all participants. This entailed the participants listening to their own recorded performance following each trial. In the second training session, the participants were divided into two groups: the Audio FB, and Augmented FB groups. The Audio FB group continued with the audio FB training, while the Augmented FB group received both sound of their own recorded performance and the difference in the waveforms of the key movement as feedback.

The experiment consisted of three test sessions (pretest, midtest, and posttest), with two training sessions between the two of these three test sessions. During the test sessions, participants listened to the recorded sound of the target performance and were asked to reproduce it five times for each test session (Figure 2B). In the first training session (Training 1), all training groups performed auditory feedback training, where they listened to their own recorded performance after each trial. This training method simulates the common piano practice, in which pianists review their own playing by listening to the recorded sound. In the second training session (Training 2), the two groups underwent different training conditions. Audio FB group continued the auditory feedback training (i.e. the same as the Training 1). Augmented FB group received both visual feedback on the key movement differences between the target and the participantʼs performance and auditory feedback of their own performance. The differences in the waveforms of the piano key movements were displayed on a monitor placed beside the piano. Before the training sessions, the participants engaged in a 5-minute familiarization period. During the familiarization period, the participants experimented with various approaches to playing, and following each trial, an experimenter explained the feedback information provided from the system to facilitate the participants’ understanding of the relationship between their performance and the feedback. Each training session consisted of 80 trials, which were split into four blocks (i.e. 20 trials for each block), with a 1-minute break between the blocks. Participants had 15 seconds to review the performance feedback after each trial.

Effectiveness of the training was assessed by the improvement of average cosine similarity relative to the pretest. The improvement was quantified as follows.

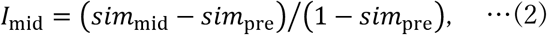

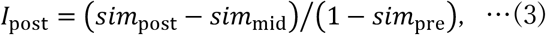

where *I*mid, and *I*post represent the improvement of the key movements similarity at the mid-test, and the post-test, respectively, whereas *sim*pre, *sim*mid, and *sim*post represent the average cosine similarity of the key movements at the post-test, mid-test, and pre-test, respectively. The denominator (1-*sim*pre) is indicative of the potential for enhancement in similarity from the pre-training state. Here, 95% confidence intervals were computed to assess the statistical significance of the improvement.

#### Analysis of Movement Exploration

We examined the extent of exploration of the movements during each feedback training session. The velocity of the key movements were calculated using central differencing of the measured displacement. The Dynamic Time Warping (DTW) with Euclidean distance was calculated between all pairs of 80 training trials, generating an 80×80 distance matrix. Multidimensional scaling (MDS) was applied to the distance matrix to project the key movement data onto a two-dimensional space. The area of the alpha-shape encompassing all training trials in this space was defined as the amount of exploration of the movements. An alpha shape is a computational method that estimates the polygonal boundary that encloses a set of points in a plane. The computational process commences with the construction of a Delaunay triangulation of the data points projected onto a plane. This triangulation method partitions the point set into triangles that are as geometrically balanced as possible. To mitigate the influence of outliers, a hyperparameter α is specified. In the triangulation process, any triangle whose circumcircle radius exceeds the α is excluded from further consideration. The remaining triangles are then connected to form the alpha shape (Edelsbrunner et al., 1983). In the present study, the value of α was defined as the mean Euclidean distance to the five nearest neighbors for all points. A larger area of alpha shape suggests that the points are more dispersed, indicating a greater variety of piano key movements during training sessions (see Figure 4A). Increment of the movement exploration was determined as the ratio between the extent of movement exploration during the first and second training sessions. A t-test was performed to determine whether the increment of exploration between the two training sessions differed significantly across the groups.

**Figure 3.**
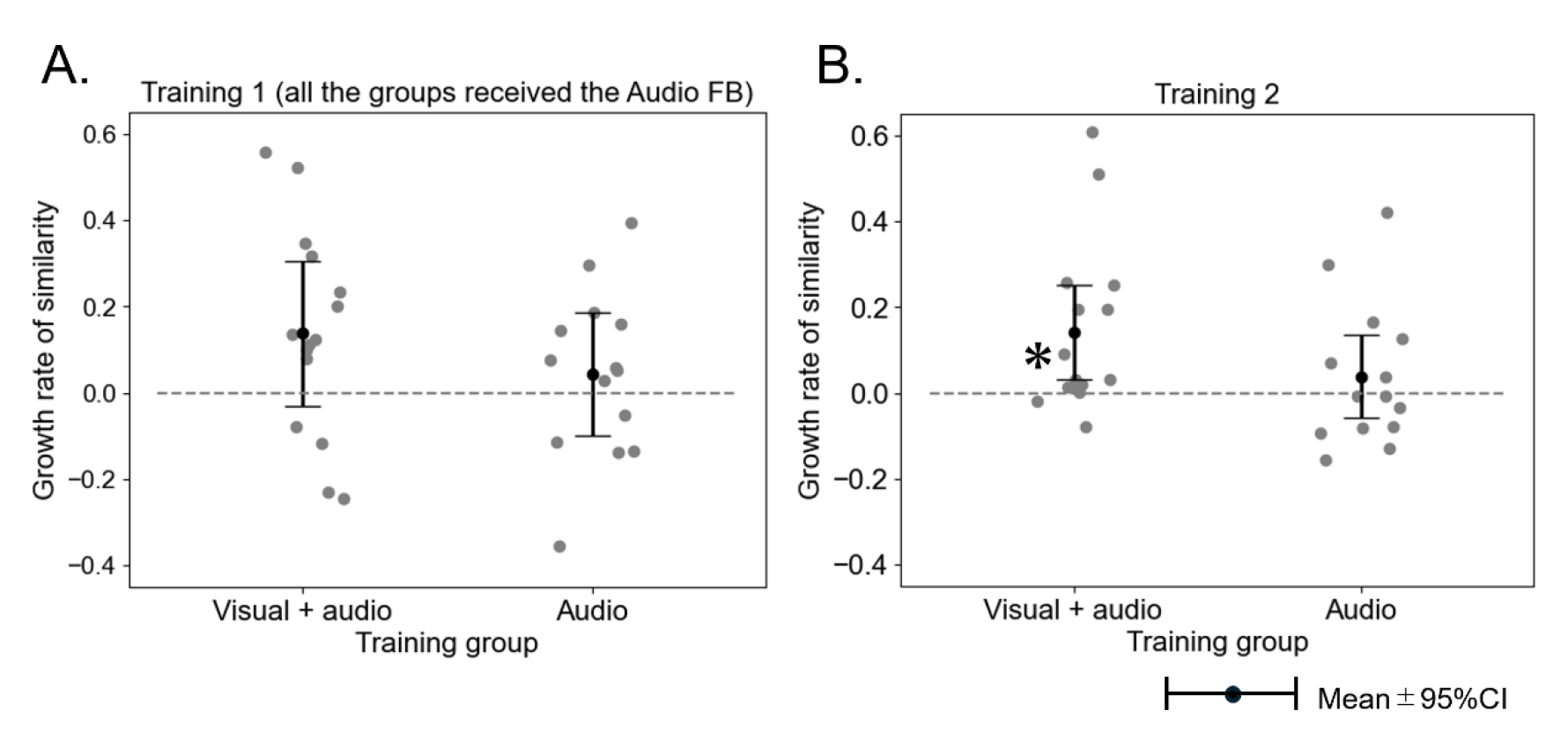
The growth rate of similarity of the key movements through Training 1 and Training 2 sessions. A: During the first training session, during which all groups underwent Audio FB training, no significant improvement in similarity of the key movement was exhibited by any of the groups. B: In contrast, during the later training session, only the Augmented FB group demonstrated a significant improvement in similarity of the key movement.

**Figure 4.**
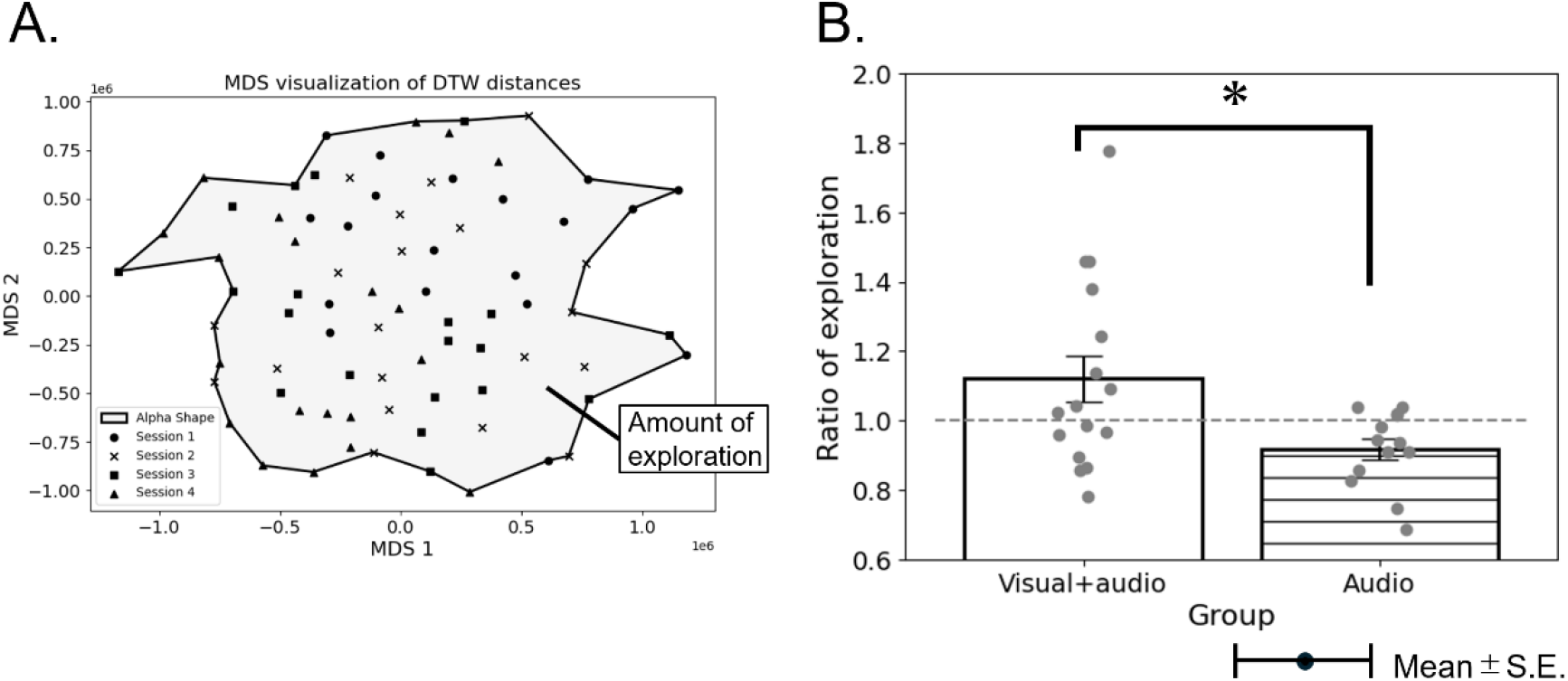
The increased ratio in the extent of exploration of movement between the first and the second training sessions. A: A representation of the projection of the key movement data during the training session onto the two-dimensional MDS space, with the alpha shape enclosing all trials. The extent of exploration of movement is defined as the area of the alpha shape that encloses all the points projected onto a two-dimensional plane using multidimensional scaling from the DTW distance matrix of each training trial. B: The augmented FB training group exhibited a substantially greater increase in the extent of movement exploration compared to the audio FB training group.

### Experiment 2: Changes in Perceived Similarity of Sound

In Experiment 2, we investigated whether feedback training improved the auditory similarity of performances through performing a listening test. In total, 46 pianists majoring in piano performance at music conservatories participated in this listening experiment. None of the listeners had participated in Experiment 1. Each participant listened to the recordings of all the participants in the Audio FB and the Augmented FB group at the mid-test and post-test sessions. In each listening trial, listeners first listened to the audio recording of the target performance as a reference, and then listened to two recordings of the same pianist, one of which was the mid-test recording and the other was the post-test recording. The listeners were asked to judge which of the two performances more resembled the reference performance. The order of providing the mid-test and post-test recordings was randomized. The training effect was defined as the percentage of pianists whose post-test performance was judged more similar to the target performance compared with the mid-test performance. A paired t-test was used to assess the difference in the training effect between the Audio FB and the Augmented FB groups.

## Results

### Changes in Key Movement Similarity through Training

Figure 3 illustrates the improvement in the similarity of key movements to the target performance through the Training 1 and Training 2 sessions, displaying the group mean of the growth rate in similarity. In the Training 1 session, the improvement score (*I*mid) for the augmented FB group and the audio FB group were 0.130±0.060 (mean ± standard error (SE)) and 0.043±0.050, respectively. Here, 95 % confidence intervals (CI) of the growth rate of the similarity were estimated to test whether the improvement was significant. The resulting confidence intervals were [-0.031, 0.305] for the augmented FB group, and [-0.099, 0.185] for the Audio FB group. As both intervals included zero, no significant improvement in similarity was observed in both groups during the first training session (Figure 3A). On Training 2 session, the improvement scores (*I*post) for the augmented FB, and audio FB groups were 0.141±0.049, and 0.038±0.043respectively. The corresponding 95% CIs were [0.031, 0.231], and [-0.058, 0.134] for the aforementioned training groups. Only the Augmented FB groupʼs CIs were above zero, indicating a statistically significant improvement in key movement similarity during the second training session (Figure 3B).

### Exploration of movements through training

Figure 4A illustrates the example of the spatial distribution of the projected key movement data in the two-dimensional MDS space, with the alpha shape enclosing all trials. The area of alpha shape indicates the amount of movement exploration. Figure 4B illustrates the changes in the amount of movement exploration through the second training session, displaying the group mean of the increase ratio of the movement exploration for each of the training groups. The growth ratio of the movement exploration between the first and second training sessions for the augmented FB, and audio FB groups were 1.120±0.066 (mean ± SE), and 0.916±0.030, respectively. A t-test indicated that the Augmented FB group exhibited significantly greater movement exploration than the audio FB group (*p*=0.018). The results indicate facilitation of movement exploration specifically through the augmented FB training.

### Changes in Similarity of Perceived Sounds

To confirm that the changes in the key movement waveforms through training accompanied perceptual changes in the tone sequences, the listening test was performed. Figure 5 illustrates the changes in the perceived similarity of the tone sequences following the second training session for the augmented FB and the audio FB group. In the augmented FB group, the percentage of the number of pianists whose sounds were judged to be more similar to the target sound following the second training session was 53.2 ± 1.8 % (mean ± standard error), while in the audio FB group, the ratio was 44.0 ± 1.9 %. A paired t-test revealed that this ratio was significantly greater in the augmented FB group than in the audio FB group (t(45) = 3.184, p = 0.002). The result indicates that multimodal FB training enhances not only the similarity of the key movements but also the perceived similarity of produced sounds to the target performance.

**Figure 5.**
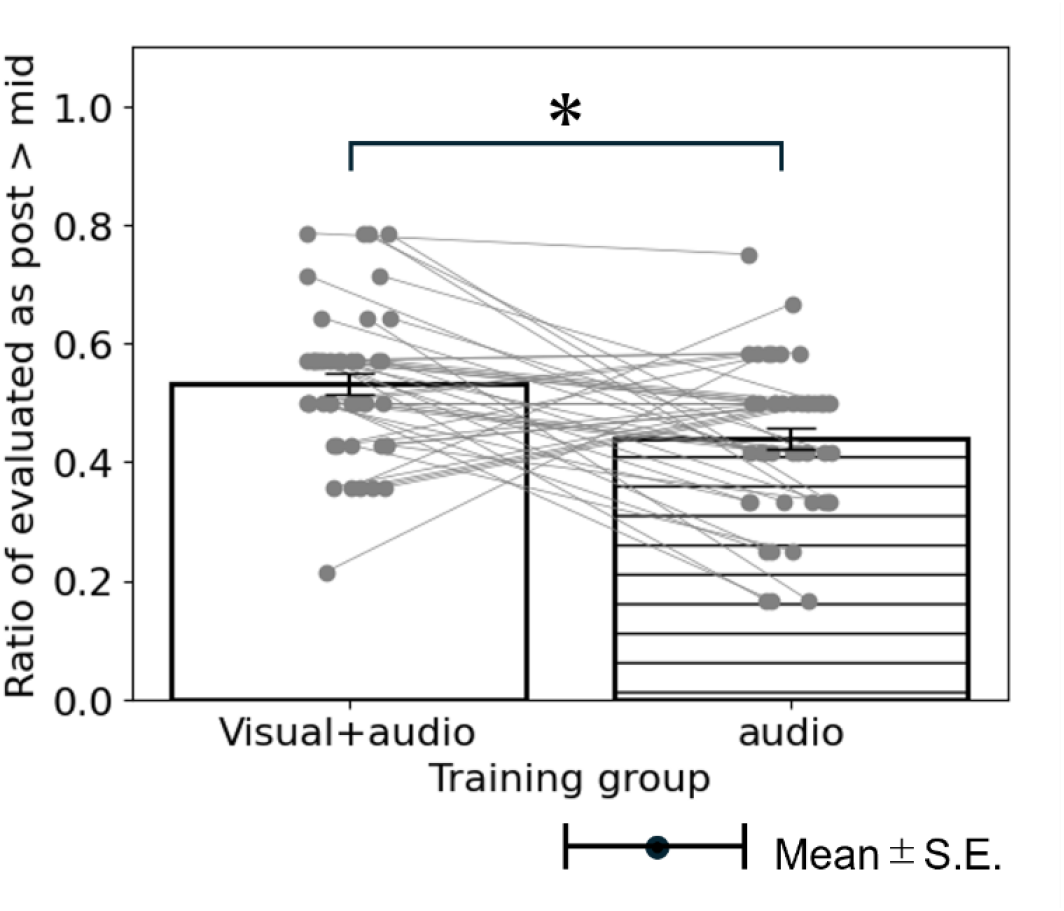
The changes in perceived similarity of sound following the second training session. The ratio of the pianists whose sounds were judged to be more similar to the target sound following the second training session was significantly greater in the Augmented FB training group than in the Audio FB training group.

## Discussion

In this study, we investigated augmented feedback grounded in high-precision sensing system can refine the performance of trained pianists, a population often considered resistant to further improvement of motor skills. By visualizing subtle differences in key motion trajectories between the learner and an expert target, this training pipeline enabled participants to achieve closer alignment with the expert performance than was possible with the conventional training with auditory feedback alone. Importantly, despite the complex mechanics of a grand piano, our listening tests with pianist participants confirmed that the augmented feedback also translated into perceptible gains in auditory similarity to the expert performance. This indicates that the benefits of feedback extended beyond mechanical mimicry to improvements with real perceptual and artistic salience. Furthermore, the augmented feedback induced a larger degree of movement exploration during training, suggesting that exposing pianists to precise, trial-by-trial discrepancies can facilitate performance refinement, with destabilizing habitual motor patterns and exploring alternative solutions contributing to the increased learning gains. Taken together, these results highlight a novel pathway for breaking expert plateaus: rather than reinforcing existing routines, sensor-mediated feedback expands the exploratory space of skilled action and drives convergence toward higher-level performance.

In this experimental design, both groups first underwent an auditory feedback training phase prior to being assigned to either the augmented feedback or auditory feedback condition for the second phase. The absence of performance enhancement during the initial auditory feedback phase suggests the presence of a ceiling effect (Furuya et al., 2014, 2025; Hirano, Sakurada, et al., 2020), indicating the participants’ expertise as pianists. For such expert performers, auditory feedback alone was not sufficient for reducing performance discrepancies relative to the target expertise, possibly because these differences are not easily perceived through the performerʼs auditory system.

Previous research has shown that simple visual feedback on performance outcomes or movement often yields diminishing returns for experts, while being more beneficial during the early phases of complex motor learning in novices (Hirano, Sakurada, et al., 2020). This contrasting result can reflect fundamental differences in the state of the learners; experts possess a highly tuned internal model of sensorimotor control (Hirano, Kimoto, et al., 2020; van der Steen et al., 2014) and exhibit enhanced sensory acuity (Evans et al., 2013; Han et al., 2015; Ragert et al., 2004; Rammsayer et al., 2012). Such advanced representation enables experts to detect performance errors without the aid of external feedback. However, our study demonstrated that when the feedback surpasses the intrinsic limits of human perception, by reaching spatial resolutions of 0.01 mm and temporal resolutions of 1 ms used in this study (Borngräber et al., 2022; Friberg & Sundberg, 1995; Van Boven & Johnson, 1994), it can detect discrepancies that would be inaccessible with the sensory system itself. By extracting keystroke-related motion features from multidimensional time-series data and amplifying the differences from the target expertise, our system improved movement similarity, thereby establishing that high-precision error amplification can extend the trainability of experts beyond the ceiling imposed by intrinsic feedback.

The feedback interface developed in this study also contributed to the training effectiveness. Rather than overwhelming performers with continuous streams of raw data, the interface balanced global indicators of performance similarity with detailed waveform-level error visualization, while granting the user autonomy to select which notes to be attended. This design aligns with prior evidence that overly frequent or externally imposed feedback can foster dependency and disrupt automatization of skill (Krause et al., 2018; Salmon et al., 1984). By structuring feedback in a self-directed manner, the system transformed a potentially intrusive volume of data into actionable and non-disruptive information, preserving automaticity while enabling targeted correction. Such an approach highlights the importance of not only what feedback is provided, but also how it is delivered, underscoring the role of interface design in human‒machine co-adaptation.

Our results also illuminate the mechanisms by which precise error feedback promotes learning. Motor skill learning is commonly modeled as the interplay of error-based learning, which updates internal models based on error with a target (Kawato, 1999; Todorov & Jordan, 2002; Wolpert & Ghahramani, 2000), and reinforcement learning, which updates motor strategies based on reward prediction(Izawa & Shadmehr, 2011). In the early stages of a novel sensorimotor task, exploration is known to facilitate skill improvement by exposing the learner to a broader behavioral repertoire (Dhawale et al., 2017; Uehara et al., 2019). In the present experiment, the auditory feedback group may have perceived their performance as already sufficiently close to the target and thus reduced exploration. Conversely, the augmented feedback group, confronted with subtle but veridical discrepancies, continued to search for elaborated motor strategies. This suggests that high-resolution feedback does not merely sharpen error correction, but also sustains the exploratory behaviors that are essential even in advanced stages of expertise. Thus, the observed performance gains likely reflect the combined action of error-based and reinforcement-based learning, triggered by feedback that challenges the performerʼs assumption of sufficiency.

This study has several limitations. We did not examine long-term retention of the training effects, nor did we evaluate whether the benefits of high-precision feedback generalize to other motor tasks. Nevertheless, the present findings have potential applications in fields such as sports, performing arts, musical performance, and traditional craftsmanship, in which years of repetitive training is required, and skill refinement often involves imitating the movements of experts. In such contexts, where expert actions are typically rapid and precise and self-perception of posture during movement is difficult, high-precision sensing combined with augmented feedback to highlight performance errors may provide an effective avenue to extend the adaptive capacity of experts.

## Acknowledgement

This study was supported by JST CREST (JPMJCR20D4) and JST CRONOS (JPMJCS24N8).

